# Induction of anergy in human T lymphocytes by exposure to single amino acid mutated antigens

**DOI:** 10.1101/2025.04.15.649051

**Authors:** Yoko Nakai-Futatsugi

**Affiliations:** RIKEN Center for Biosystems Dynamics Research, Laboratory for Retinal Regeneration

**Keywords:** anergy, autoimmune disease, T cell, antigen, peptide

## Abstract

Anergy is a condition of immune T cells becoming sustainably inactive. Theoretically, it occurs when a T cell recognizes its antigen but cannot be active. If we could artificially induce anergy to a specific T cell clone, this should provide a therapeutic strategy for autoimmune disease. One possible method to induce anergy is to expose a T cell to an unstably presented antigen, which makes the T cell bind to the antigen but not become active due to insufficient duration of antigen presentation. The present study attempted to realize this condition in human T lymphocytes by single amino acid mutation of the antigen. As a result, the potentiality of this method for anergy-induction in human T lymphocytes was revealed. By *in silico* protein structure prediction, it was suggested that optimal range of contact energy between the antigen and the major histocompatibility complex was required for successful induction. From the gene expression profile, emergence of CD4+/TSHZ2+ T cell population was suggested as one of the hallmarks of anergy-induction by this method. This was the first study to demonstrate anergy-induction by the destabilized antigen strategy in human cells.

## Introduction

T lymphocytes are immune cells that are tailored to recognize a specific antigen. The acquired immune system driven by T lymphocytes is one of the most complicated systems in the body. To protect our bodies from upcoming pathogens during life, a huge repertoire of T cell clones is prepared, followed by selection at the thymus around birth to eliminate the clones that recognize organ-specific molecules of the same body. The selection is not too strict, and is continuously occurring even in adulthood, making the immune system responsive to a variety of pathogens. However, when the selection is dysregulated, the organs could be attacked by its own immune system. This is what is called autoimmune disease. Autoimmune disease is generally chronic and intractable. Despite the increasing incidence of autoimmune diseases, the treatment for now is by immune suppressive drugs that systemically suppress the immune system, which accompany significant adverse effects. Therefore, more safe and efficient therapeutic strategy, ideally by exclusive suppression of a specific T cell clone is desired.

Besides the aforementioned initial selection of the immune T cells at the thymus, peripheral selection in the blood also takes place during life. One of the ways of the selection to suppress self-recognizing T cells is by induction of anergy. Anergy occurs under a condition in which a T cell recognizes its antigen but cannot be active (in other words, by non-functional recognition of an antigen), leading to a selective and eternal unresponsiveness of that T cell (reviewed in Appleman and Boussiotis) [1]. If this innate selection can be induced artificially, it may be an ideal therapeutic strategy for autoimmune disease.

T cells become active when the T cell receptors (TCR) binds to antigen-peptides presented on the major histocompatibility complex (MHC) at the cell surface of antigen-presenting cells. Previous studies have demonstrated induction of anergy by blocking of the costimulatory receptor CD28 simultaneously with the binding of TCR to the antigen/MHC-complex [2]. The involvement of CD28 in anergy induction is supported by a study with a knockout mouse of cytotoxic T lymphocyte antigen 4 (CTLA4), the negative immune regulator that competes with CD28, showing the requirement of CTLA4 for induction of anergy *in vivo* [3]. The immune suppressive cytokine interleukin (IL)-10 applied simultaneously with CD4+ T cell activation also induces anergy *in vitro* [4]. Although these lines of evidence show induction of clonal anergy in T cells, it targets CD28, CTLA4 or IL10 that are general components of the immune system so theoretically any T cell clone can be affected. Alternatively, anergy in a specifically targeted T cell clone was able to be induced by making the TCR bound to an unstable antigen/MHC-complex [5]. As the duration of antigen presentation matters for T cell activation [6], unstable loading of the antigen peptide on the MHC should lead to insufficient activation of the T cell even after the TCR transiently binds to the antigen/MHC-complex. Based on this idea, Ford and Evavold [7] have made the antigen/MHC-complex destabilized by single amino acid mutation to the MHC-anchor residue of the antigen-peptide, which indeed suppressed the hyperactive immunological phenotype of autoimmune encephalomyelitis mouse model.

The present study applied the method of Ford and Evavold [5,7] to human T lymphocytes. Although it was stochastic, induction of anergy by single amino acid mutation at the MHC-anchor residue of the antigen-peptide was somehow demonstrated, suggesting the therapeutic potential of this method for autoimmune disease.

## Results and Discussion

### In vitro induction of anergy by single amino acid mutated antigen-peptide

The amino acid sequence of an antigen loaded on an MHC molecule is specific to the subtype of the MHC. In this study, cytomegalovirus (CMV)-antigen loaded on the A24 subtype of human leukocyte antigen (HLA; the MHC molecule of human) was used as a model of T cell recognition of MHC/antigen-complex. CMV-antigen was chosen because most of the human beings are potentially infected by CMV so blood samples with T cell clones that are active against CMV-antigen is generally available, and the antigen sequences of CMV for each HLA subtype are well characterized. The amino acid sequence of CMV-antigen loaded on HLAA24 is known as QYDPVAALF. Following the method of Ford and Evavold [5,7], mutation was made to the MHC-anchor residues (PVA) by replacing the hydrophobic residues P and A with hydrophilic residues K and S, respectively, resulting in two single mutated peptides (QYD<u>K</u>VAALF and QYDPV<u>S</u>ALF) and one double mutated peptide (QYD<u>K</u>V<u>S</u>ALF) for test. First, whether these mutations affect the binding to HLAA24 was assessed by *in vitro* competition assay (**Fig. 1A**). HLAA24-monomer coated on a dish was incubated with biotinylated antigen (QYDPVAALF-biotin) together with the original (QYDPVAALF) or mutated (QYD<u>K</u>VAALF, QYDPV<u>S</u>ALF or QYD<u>K</u>V<u>S</u>ALF) antigen-peptides, then the amount of QYDPVAALF-biotin bound to the coated HLAA24-monomer was measured. The mutated peptides were less competitive compared to the original antigen, indicating the binding to the HLA was efficiently weakened by single amino acid mutation of the antigen (**Fig. 1A**).

**Fig. 1.**
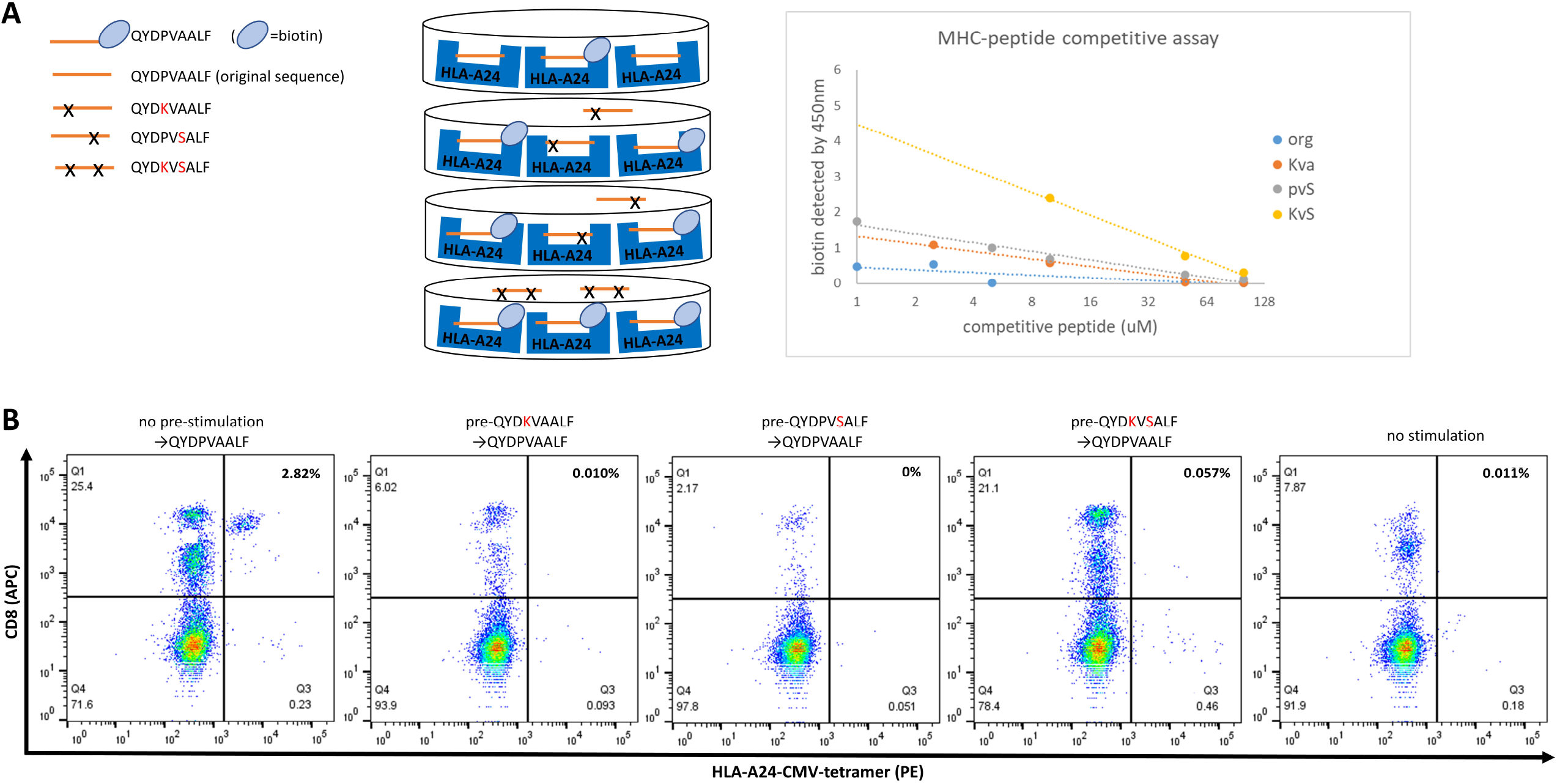
Single amino acid mutated antigen with less affinity to the HLA attenuated the T cell clone. **A** HLAA24-monomer coated on a dish was incubated with biotinylated antigen together with mutated antigens. The amount of biotinylated antigen bound to the HLAA24-monomer in competition with the original antigen or mutated peptides was measured. Mutation by single amino acid gave ambiguously lowered affinity. **B** HLAA24-positive PBMC obtained from a healthy doner (HD1) were incubated with mutated peptides, followed by the stimulation with the original CMV-antigen. HLAA24-CMV-tetramer positive area indicates T-cell clones that recognize HLAA24/CMV-antigen-complex. The number on the upper right of each plot indicates the percentage of HLAA24-CMV-tetramer positive/CD8+ T cell population.

Next, the effect of ambiguously weakened binding between HLA and antigen onto the activity of HLAA24/CMV-responding T cell clone was tested. Peripheral blood mononuclear cells (PBMC) were obtained from a HLAA24-positive healthy doner (HD1). HLAA24-CMV tetramer (MBL Life Science) was used to detect the T cell clone that recognized HLAA24/CMV-complex upon stimulation of PBMC with the original CMV antigen (**Fig. 1B**, most left plot). When the PBMC was pre-stimulated with single amino acid mutated peptides for 24 hours, followed by incubation with the original CMV antigen for 2 weeks, HLAA24/CMV-responding T cells were not detected (**Fig. 1B**, 2nd and 3rd plots), which was similar to the condition without any stimulation with the CMV-antigen (**Fig. 1B**, most right plot), suggesting elimination or attenuation of the HLAA24/CMV-responding T cell clone by 24-hour exposure to single amino acid mutated CMV-antigens. When double amino acid mutated CMV-antigen was used for the 24-hour pre-stimulation, CD8+ T cells other than HLAA24/CMV-responders, presumably responders to the antigen closer to the double mutated sequence, were activated (**Fig. 1B**, the plot 2nd from the right). These results indicate that a specific T cell clone is able to be inactivated by exposure to ambiguously unstable HLA/antigen-complex, which may represent induction of anergy.

### Stochastic feature of anergy-induction by single amino acid mutated antigens

The same experiment of **Fig. 1B** was repeated in duplicated wells of a 96 well plate, for two HLAA24-positivie healthy doners (HD1 and HD2)-derived PBMCs. The population of CD8+ T cell that recognized HLA/CMV-complex (HLAA24-CMV-tetramer positive/CD8+ population) was quantified (**Fig. 2, Fig. S1)**. For HD1, one of the two wells stimulated with QYDPV<u>S</u>ALF mutated antigen did not show successful attenuation of the T cell clone (**Fig. 2, Fig. S1**), indicating even the same batch of PMBC samples exposed to the same mutated antigen under the same experimental condition would give different results probably due to the microenvironment of the PBMC culture. For HD2, attenuation to the T cell clone was induced less efficiently, as decrease in HLAA24-CMV-tetramer positive population was shown in none of the two wells by QYD<u>K</u>VAALF, and in one of the two wells by QYDPV<u>S</u>ALF (**Fig. 2, Fig. S1**). So, the mutated antigen QYD<u>K</u>VAALF induced anergy-like attenuation of the T cell clone in HD1 but not in HD2, while the mutated antigen QYDPV<u>S</u>ALF induced anergy-like attenuation both in HD1 and HD2, although it occurred only in half of the wells for both doners (**Fig. 2, Fig. S1**). It could be speculated that not too much but an ambiguous instability of the antigen/HLA-complex is required for successful anergy-induction, because if the binding between HLA and antigen is too low, the antigen will not be loaded on the HLA so will not be recognized by the TCR; and if the binding is too stable, it will be recognized by the TCR as normal antigen so the T cell clone will be even more activated. This ambiguousness of peptide design and stochasticity of anergy induction could be the reasons why this apparently simple and safe method has not yet been clinically applied.

**Fig. 2.**
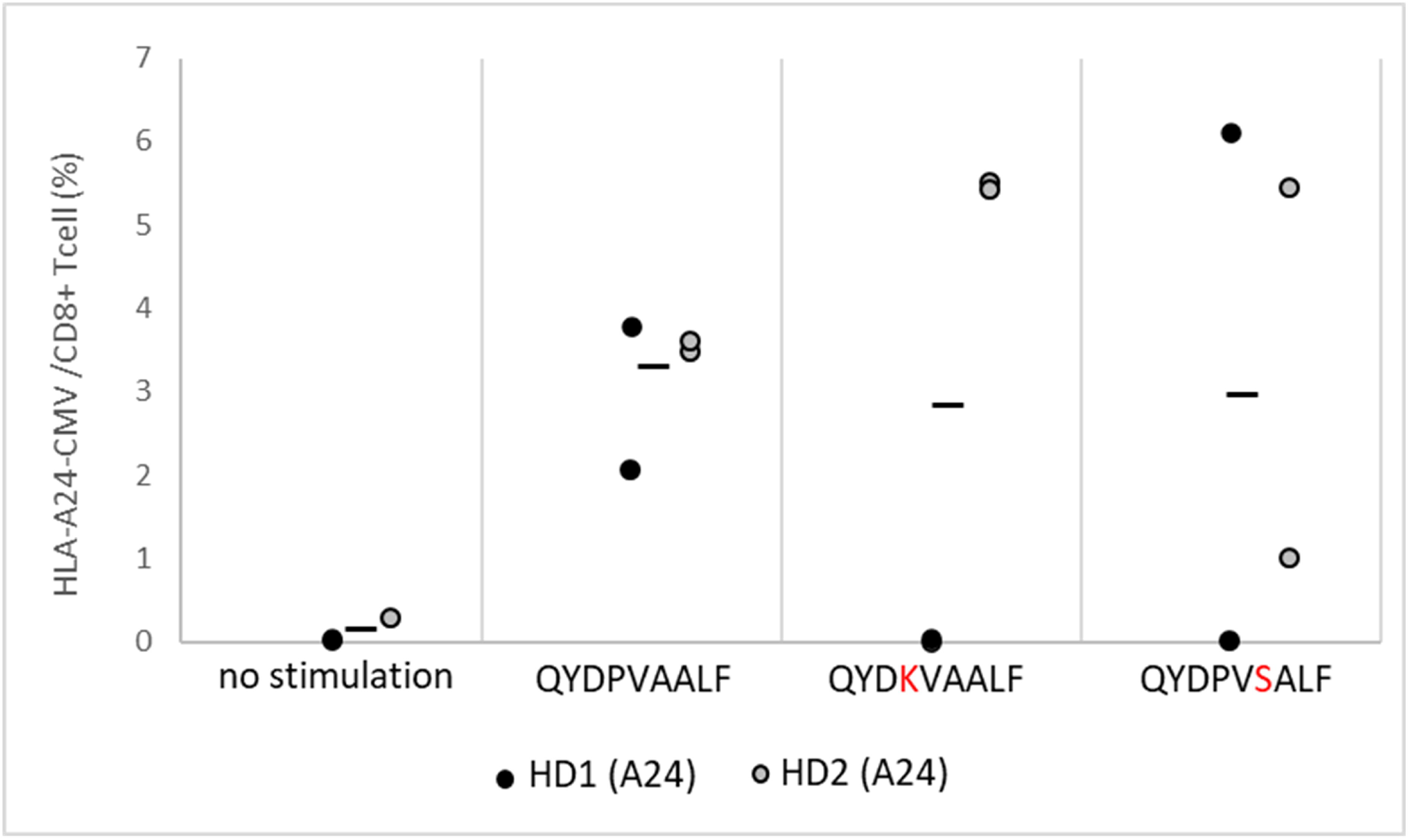
Induction of anergy by single amino acid mutated antigens was stochastic. The experiment of **Fig. 1B** was repeated in duplicated wells for PBMCs derived from two HLAA24-positive healthy doners (HD1 and HD2), from which HLAA24-CMV-tetramer positive/CD8+ population was quantified. The actual plots are in **Fig. S1**. Y-axis indicates HLAA24-CMV-tetramer positive/CD8+ population (%). Each dot represents HLAA24-CMV-tetramer positive/CD8+ population in each plot of **Fig. S1**. Black line indicates average.

### In silico analysis of HLA-antigen binding

The binding efficiencies between HLA and mutated antigens were predicted with a protein structure prediction software, Molecular Operating Environment (MOE; Chemical Computing Group) with the HLABAP module [8] (MOLSIS Inc.) (**Fig. 3**).

**Fig. 3.**
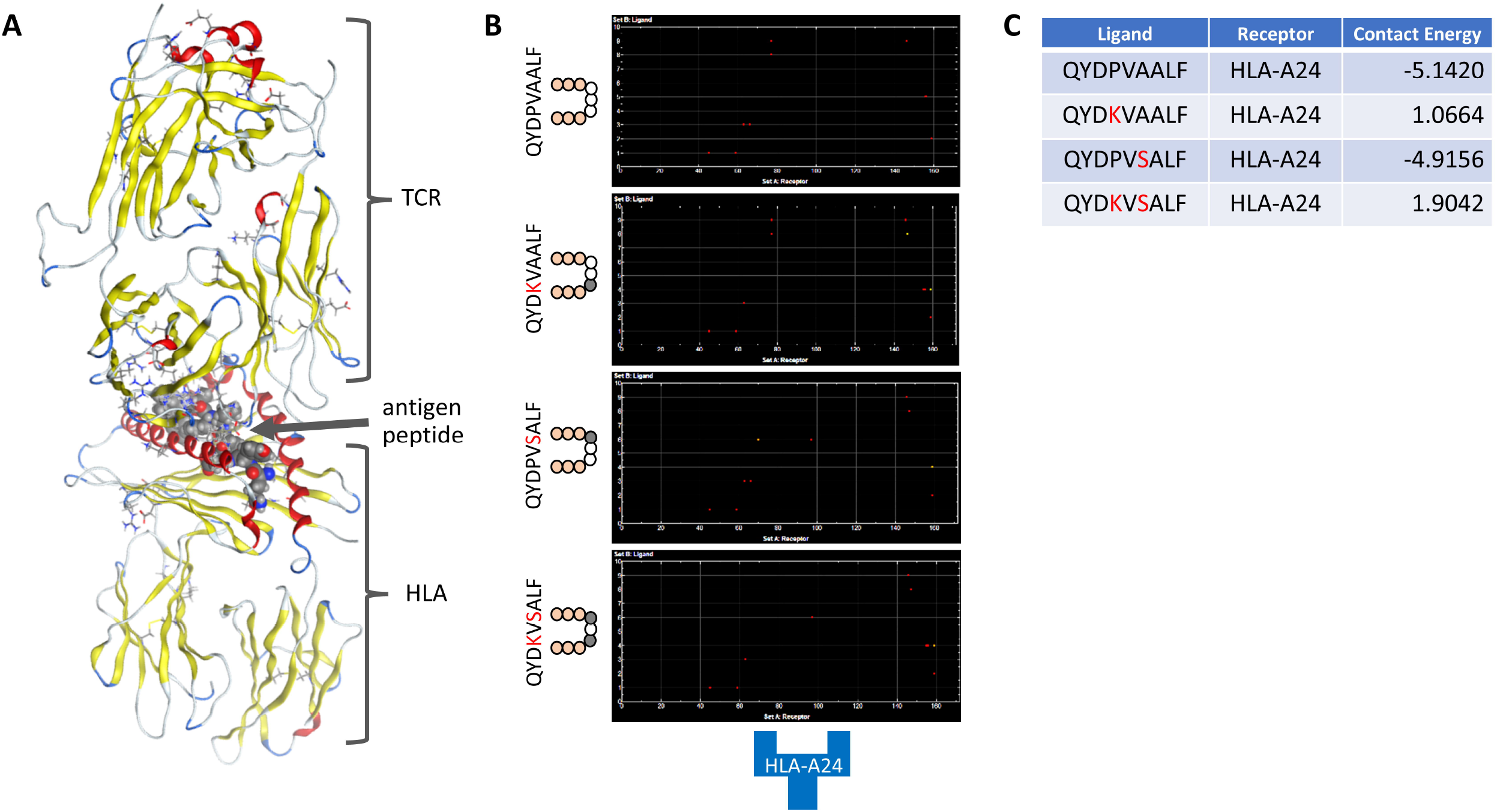
Peptides with single amino acid mutation had lower affinity to HLA. **A** Protein structure of peptide antigen bound to the pocket of HLA and recognized by the TCR was depicted with the MOE software. **B** The binding positions between antigen peptide (y-axis) and HLA (x-axis) at amino acid level were predicted with the MOE software. Red and orange dots indicate predicted hydrogen bonds and arene bindings, respectively. **C** Affinity of antigen peptide to HLA was predicted with the MOE-HLABAP software. Lower contact energy indicates higher affinity. Mutation by single amino acid gave lowered affinity that differed depending on the mutated residue, which was much lowered by double amino acid mutation.

The binding position of the residues within the complex were slightly altered by amino acid mutation of the antigen (**Fig. 3B**). Among the four antigen sequences tested in **Fig. 1**, indeed the original antigen QYDPVAALF showed the highest affinity to HLAA24 with less contact energy; single amino acid mutation gave less affinity with higher contact energy; and double amino acid mutation gave the lowest affinity with the highest contact energy (**Fig. 3C**), which were consistent with the *in vitro* binding assay shown in **Fig. 1A**. Although both of the single amino acid mutated antigens (QYD<u>K</u>VAALF and QYDPV<u>S</u>ALF) showed similar binding affinity to the HLA by the *in vitro* binding assay (**Fig. 1A**), the difference between the two mutations was more evident by protein structure prediction, having HLA-contact energy of QYDPV<u>S</u>ALF (−4.9156) much closer to the original QYDPVAALF (−5.1420) compared to QYD<u>K</u>VAALF (1.0664) (**Fig. 3C**). Since QYDPV<u>S</u>ALF but not QYD<u>K</u>VAALF was able to induce anergy-like attenuation of T cell in both HD1 and HD2 derived PBMCs regardless of doner-specificity, and QYDPV<u>S</u>ALF was predicted to have lower HLA-contact energy than QYD<u>K</u>VAALF, it is possible that higher contact energy to the HLA (*i*.*e*. less affinity) after single amino acid mutation of the antigen is not the determinant for anergy-induction. This notion was supported by the same experiment using HLAA2-positive PBMCs stimulated with four types of single amino acid mutated CMV-antigens (**Fig. S2**). The peptide sequence of the CMV-antigen for HLAA2 is known as NLVPMVATV. Four single mutated antigens (NLV<u>K</u>MVATV, NLVPM<u>D</u>ATV, NLV<u>R</u>MVATV and NLVPM<u>K</u>ATV) that had much higher HLA-contact energy (1.1716, 1.16281, 2.4966 and 5.8791, respectively) were tested, but successful anergy-like T cell attenuation was not induced (**Fig. S2**). As discussed earlier, it is possible that if the affinity between antigen and HLA is too low, the antigen will not be presented on the HLA during the 24-hour pre-stimulation period thus anergy will not be induced. Probably ambiguous instability at optimal range is required, for example as the slight increase in contact energy shown by HLAA24-CMV-QYDPV<u>S</u>ALF mutation (−4.9156 vs original -5.1420; **Fig. 3C**).

### Immunosuppressive effect of mutated peptide revealed by single cell analysis

To explore whether anergic profile in terms of gene expression was induced by this seemingly ambiguous and stochastic strategy, PBMC exposed to the mutated antigen peptide was analyzed by single RNA sequencing analysis. For this, HLAA24-positive HD2 PBMC stimulated with mutated QYDPV<u>S</u>ALF for 24 hours followed by stimulation with the original QYDPVAALF for 2 weeks in a 96-well plate (the condition same as the cells used in **Fig. 1B**) was prepared. QYDPV<u>S</u>ALF-mutation was used for pre-stimulation because it successfully induced anergy-like T cell attenuation in both HD1 and HD2 (**Fig. 2, Fig. S1**). Considering the stochastic effect of culture microenvironment (**Fig. 2, Fig. S1**), 5 wells from the 96-well plate were combined (total 1,000,000 cells). For control, the same batch of culture (5 wells; total 1,000,000 cells) without 24-hour pre-stimulation by mutated QYDPV<u>S</u>ALF was used. The gene expression profile of each cell obtained from RNA sequencing was divided in 20 clusters by t-distributed Stochastic Neighbor Embedding (t-SNE), of which clusters 3, 5 15, 20 (CD8+ T), clusters 0, 2, 4, 9, 13 (Memory CD4+ T) and cluster 14 (Naïve CD4+ T) were T cell population (**Fig. 4A, Fig. S3**). Among the 20 clusters, the formation of cluster 3 (CD8+ T; **Fig. 4A** red circle) and the cell number of cluster 0 (Memory CD4+ T; **Fig. 4A** black circle) were apparently different between control and QYDPV<u>S</u>ALF-exposed. When cluster 3 was divided in 8 subclusters (**Fig. 4B**), cluster 3.3 (**Fig. 4B** red circle) and cluster 3.8 (**Fig. 4B** red dashed circle) were exclusively populated in QYDPV<u>S</u>ALF-exposed and the control, respectively. The most differentially expressed marker of cluster 3.3 (the cluster exclusively populated in QYDPV<u>S</u>ALF-exposed) was a non-coding RNA AL606807.1 (**Table S1**), which was also expressed in cluster 3.8 (the cluster exclusively populated in the control) (**Fig. 4C, Fig. S4**). Both clusters 3.3 and 3.8 predominantly expressed CCL3 and CCL4 (**Fig. 4C, Fig. S4**) that are known to be expressed by T regulatory cells (Tregs) to suppress CD4+ and CD8+ T cells [9], suggesting the immune suppressive profiles of clusters 3.3 and 3.8. However, PDCD1 (also referred to as PD-1) that is expressed by antigen-experienced CD8+ T cells leading to CD8+ T cell exhaustion [10] was markedly expressed in cluster 3.3, but not in cluster 3.8 (**Fig. 4C, Fig. S4**), suggesting exposure to mutated QYDPV<u>S</u>ALF-antigen had generated a different mode of immune suppression that was not detected by stimulation with the original antigen QYDPVAALF without pre-exposure to the mutated antigen.

**Fig. 4.**
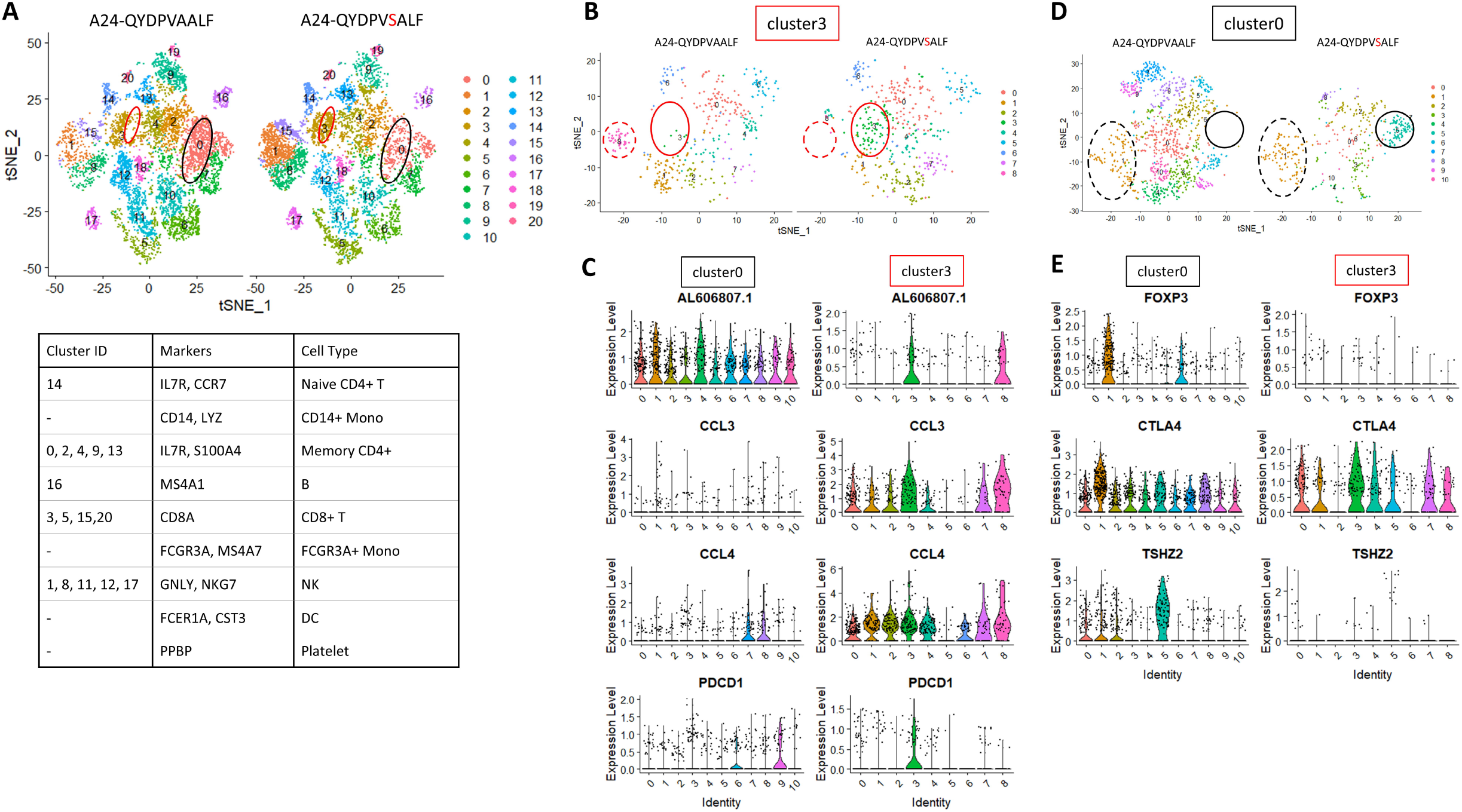
Single cell analysis of HLA-A24 PBMC stimulated with mutated CMV-antigen. **A** HLAA24 PBMC stimulated with the original CMV antigen (QYDPVAALF) or mutated antigen (QYDPV<u>S</u>ALF) were subjected to single cell RNA sequencing. When the gene expression profile was divided in 20 clusters, cluster 3 (CD8+ T, red circle) and cluster 0 (Memory CD4+ T, black circle) showed different population between the two conditions. **B** When cluster 3 was divided in 8 subclusters, cluster 3.3 (red circle) and cluster 3.8 (red dashed circle) were exclusively populated in mutated and original condition, respectively. **C** Violin plot of the subclusters of cluster 0 and 3 for the representative genes of cluster 3. PDCD1 was markedly expressed in subcluster 3.3. **D** When cluster 0 was divided in 10 subclusters, cluster 0.5 (black circle) was exclusively populated in the mutated condition, while all other subclusters except for FOXP3+ cluster 0.1 (black dashed circle) were decreased. **E** Violin plot of the subclusters of cluster 0 and 3 for the representative genes of cluster 0. TSHZ2 was markedly expressed in subcluster 0.5.

When cluster 0 (**Fig. 4A** black circle) was divided in 10 subclusters (**Fig. 4D**), cluster 0.5 (**Fig. 4D** black circle) was exclusively populated in the mutated condition, while all other subclusters except for cluster 0.1 (**Fig. 4D** black dashed circle) that expressed the Treg marker FOXP3 (**Fig. 4E, Fig. S5**) were decreased. Interestingly, cluster 0.5 that was exclusively populated by pre-exposure to mutated QYDPV<u>S</u>ALF-antigen showed marked expression of TSHZ2 (**Fig. 4E, Fig. S5, Table S2**). As TSHZ2 is implied to have an inhibitory effect on CD4+ T cell proliferation [11], the result suggests pre-exposure to mutated QYDPV<u>S</u>ALF-antigen have led to decrease in CD4+ T cells except for FOXP3+ Tregs, which have accompanied the emergence of less proliferating TSHZ2+/CD4+ T cell population instead. The TSHZ2+ cluster 0.5 did not express FOXP3 (**Fig. 4E, Fig. S5**), which was consistent with previously defined anergic T cell-profile [12]. Although the mutated CMV-antigen was supposed to affect CD8+ T cells that recognize class I HLA molecules (HLA-A, B and C, including HLA-A24 and HLA-A2 used in this study), CD4+ T cells that recognize class II HLA molecules were affected. This could be interpreted at least in part as the effect of CCL3 and CCL4 chemokines that suppress CD8+ and CD4+ T cells, which was predominantly expressed in CD8+ cluster 3.3 in response to the exposure to mutated QYDPV<u>S</u>ALF-antigen.

In summary, although not in a promising efficiency, exposure to mutated antigen that had ambiguously lowered affinity to HLA led to the emergence of anergic T cell population. *In silico* protein structure analysis may help designing an optimal affinity between HLA and antigen for anergy induction. While a well-characterized CMV antigen that are generally infected in healthy individuals was used as a model in this study, applying the strategy to real-world autoimmune diseases would have many hurdles. For example, the causal antigen should vary among patients even with same target disease, so precise sequence of the antigen has to be determined for each patient to design optimal mutation. How to deliver the peptides into human body for clinical use is also an issue. Even if anergy is efficiently induced to a specific T cell clone by the strategy, it is uncertain how long the efficacy will last systemically. Despite all the remaining uncertainty, the potentiality of this anergy-inducing strategy on human T lymphocytes to realize a safe and theoretically an eternal cure for autoimmune disease was demonstrated.

## Materials and Methods

### Cell culture

HLAA24-positive PBMCs were isolated from blood samples obtained from healthy doners (HD1 and HD2). The use of human blood sample was approved by Institutional Review Board of RIKEN Center for Biosystems Dynamics Research. HLAA2-positive PBMCs were purchased from Precision for Medicine (Frederick, MD) (HD3: lot #2010113909; HD4: lot #3010116197). For *in vitro* anergy induction, 20,000 cells/well of PBMC were seeded into multiple wells of a round-bottom 96-well plate in RPMI-10%FBS medium (RPMI-1640 medium supplemented with 10% fatal bovine serum (FBS; LONZA, Basel, Switzerland), 10mM HEPES (Sigma-Aldrich, St. Louis, MO), 0.1mM nonessential amino acids (Sigma-Aldrich), 1mM sodium pyruvate (Sigma-Aldrich), penicillin-streptomycin (Thermo Fisher Scientific, Waltham, MA), and 10μM 2-mercaptoethanol (Sigma-Aldrich)) with or without mutated antigen peptide for 24 hours. After 24 hours the medium was removed and replaced with fresh RPMI-10%FBS medium added with 10μM of the original antigen peptide. From the following day, half of the medium was replaced every other day with fresh RPMI-10%FBS medium supplemented with human-recombinant IL-2 (100 U/mL; BD Biosciences, San Jose, CA) and cultured for 2 weeks. The original and mutated antigen peptides were synthesized by Fujifilm-Wako Co. (Osaka, Japan) or BEX Co., Ltd (Tokyo, Japan). The peptides were solved in DMSO at the concentration of 10mM and stored at 4LJ; the working solution was made by dilution in the culture medium at the concentration of 1mM, and was used at the final concentration of 10μM.

### In vitro binding assay

All procedures were performed at room temperature if not specified. A 96-well ELISA plate was coated with streptavidin (Fujifilm-Wako) (2μg/ml in PBS with 10%BSA; 100μl/well) overnight, washed three times, then incubated with ultra violet (UV)-exchangeable biotinylated Flex-T™ HLA-A*24:02 Monomer UVX (BioLegend) (4μg/well in PBS with 10%BSA) for 20 minutes, washed three times, then 0.6μM of biotinylated reference peptide (QYDPVAALF-biotin; synthesized by Fujifilm-Wako) together with 0-300μM of the original (QYDPVAALF) or mutated (QYD<u>K</u>VAALF, QYDPV<u>S</u>ALF or QYD<u>K</u>V<u>S</u>ALF) antigen-peptides diluted in PBS with 10%BSA (100μl/well) were applied. The plate was sealed to avoid evaporation, and was placed in a UV-illuminator with an ice-pack on the lid, then illuminated from the bottom of the plate with 365nm UV for 30 minutes to remove pre-loaded UV-labile peptide from the Flex-T™ HLA monomer. Then the plate was placed in a 37□ incubator for 30 minutes to let the higher affinity peptides bind to the empty HLA monomer. After washed three times, the reference QYDPVAALF-biotin peptide bound to the HLA-A*24:02 monomer was detected with HRP-conjugated streptavidin and substrate Color reagent following the manufacturer’s protocol (R&D Systems, Minneapolis, MN), and measured with Multiskan plate reader equipped with SkanIt software (Thermofisher, Waltham, MA).

### Flow cytometry

PBMC were harvested and washed once with RPMI 2%FBS, incubated with Fc-block (Miltenyi Biotec) on ice for 15□minutes, washed once, incubated with T-Select HLA-A*24:02 CMV pp65 Tetramer-QYDPVAALF-PE (#TS-0020-1C; MBL Life Science, Tokyo, Japan) on ice for 30 minutes, then APC-conjugated anti-human CD8 (#17-0088-42; eBioscience, San Diego, CA) was added and incubated on ice for another 30 minutes, washed once, then analyzed by FACSLyric™ Flow Cytometry System (BD Bioscience, Franklin Lakes, NJ). FlowJo software (version 9) was used for data analysis.

### In silico binding analysis

For prediction of binding affinity between HLA and antigen peptide, Molecular Operating Environment (MOE; Chemical Computing Group, Quebec, Canada) with the HLABAP module [8] (MOLSIS Inc., Tokyo, Japan) was used.

### Single cell RNA sequencing analysis

Five wells of PBMC culture were combined and 1×10^6^ cells were subjected to RNA sequencing analysis. Preparation of cDNA library with Chromium GEM-X Single Cell 3’RNA-seq (10x Genomics, Pleasanton, CA) and RNA sequencing with Illumina Next-Generation Sequencer (Illumina Inc., San Diego, CA) were outsourced to KOTAI Biotechnologies, Inc. (Osaka, Japan). Obtained single cell RNA sequencing raw data were analyzed using Seurat v4 [13,14,15,16] with R v4.4.2 [17]. For cluster analyses in **Fig. 4**, the following parameters were used:

RunTSNE(reduction = “pca”, dims = 1:20)

FindNeighbors(reduction = “pca”, dims = 1:20)

FindClusters(resolution = 1.2)

## Acknowledgments

I thank Dr. Akira Futatsugi (Kobe City College of Nursing) for critical reading of the manuscript. This work was supported by JSPS KAKENHI Grant Number 21H03097 and RIKEN Incentive Research Project FY2021.

## Figure legends

**Fig. S1. Inconsistency of the results of the experiment in Fig. 1B**. 20,000 cells/well of PBMC derived from HLAA24-positive HD1 (upper 8 plots) or HD2 (lower 7 plots) were seeded into duplicated wells of a round-bottom 96 well plate and stimulated with mutated CMV antigens for 24 hours then cultured in the presence of original antigen for 2 weeks. T cell clone attenuation by mutated antigens was not consistent depending on the doner or the microenvironment of the culture.

**Fig. S2. T cell clone attenuation did not occur by HLAA2-CMV mutated antigen**. PBMC derived from two HLAA2-positive healthy doners (HD3 and HD4) were pre-stimulated with two types of single amino acid mutated antigens followed by stimulation with the original antigen, but the HLAA2-CMV-tetramer-positive T cell clone did not decrease. PBMC derived from HD3 was pre-stimulated with four types of single amino acid mutated antigens followed by stimulation with the original antigen, but the HLAA2-CMV-tetramer-positive T cell clone did not decrease. HLA-contact energy of the mutated antigens were predicted by the MOE-HLABAP software.

**Fig. S3. Cell type profiling of Fig. 4A**. Gene expression profile of PBMC was divided in 20 clusters by t-SNE. Expressions of cell type markers plotted by FeaturePlot are shown.

**Fig. S4. Cluster 3 gene expression**. Cluster 3 in Fig. 4A was subdivided in 8 clusters. Gene expressions of the markers of subcluster 3.3 plotted by FeaturePlot are shown.

**Fig. S5. Cluster 0 gene expression**. Cluster 0 in Fig. 4A was subdivided in 10 clusters. Expressions of critical genes plotted by FeaturePlot are shown. TSHZ2 was markedly expressed in cluster 0.5.

## Notes

### Competing Interest Statement

The authors have declared no competing interest.

