## Supplementary figures and images for "Induction of anergy in human T lymphocytes by exposure to single amino acid mutated antigens"

### Fig. S1

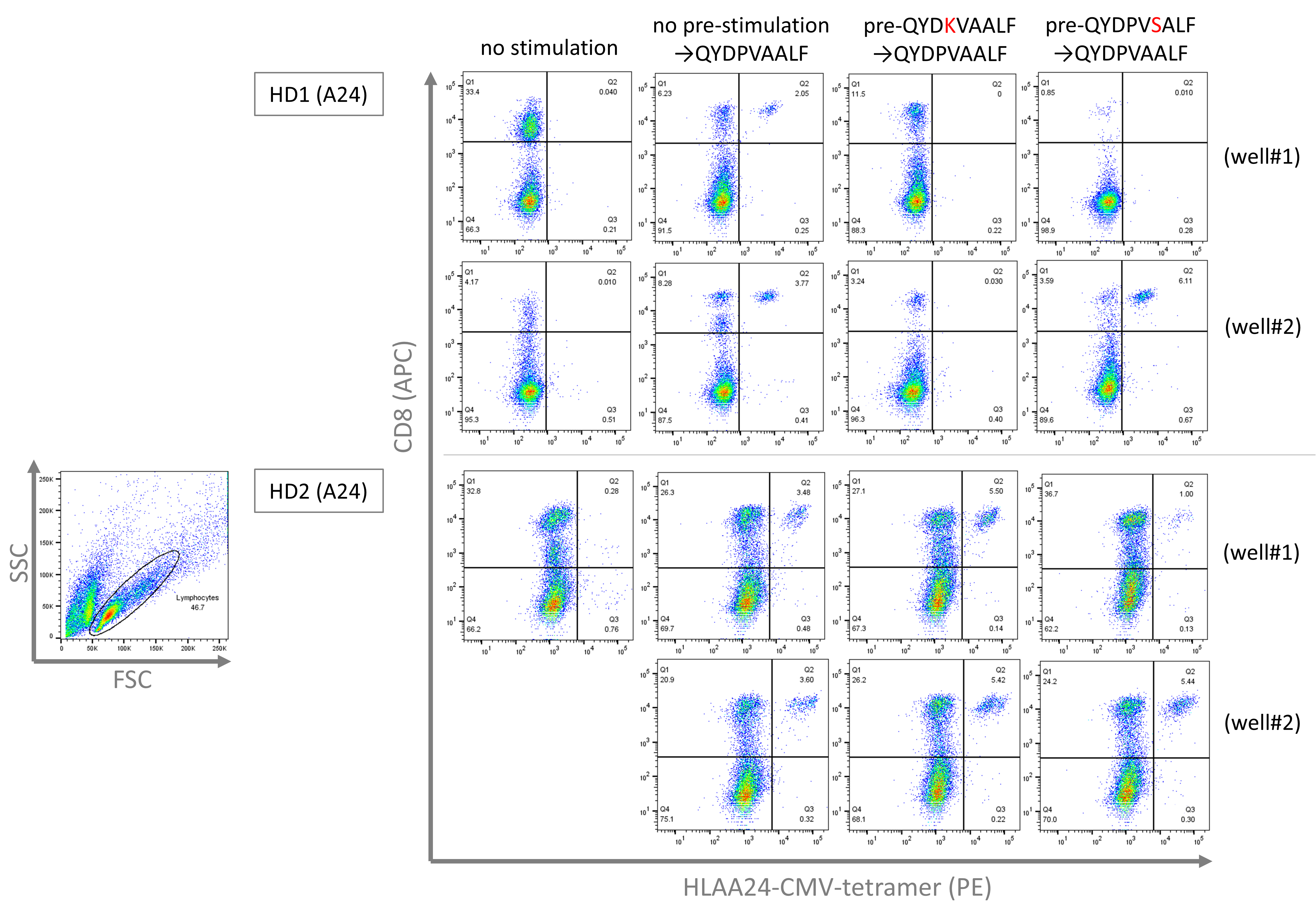

### Fig. S2

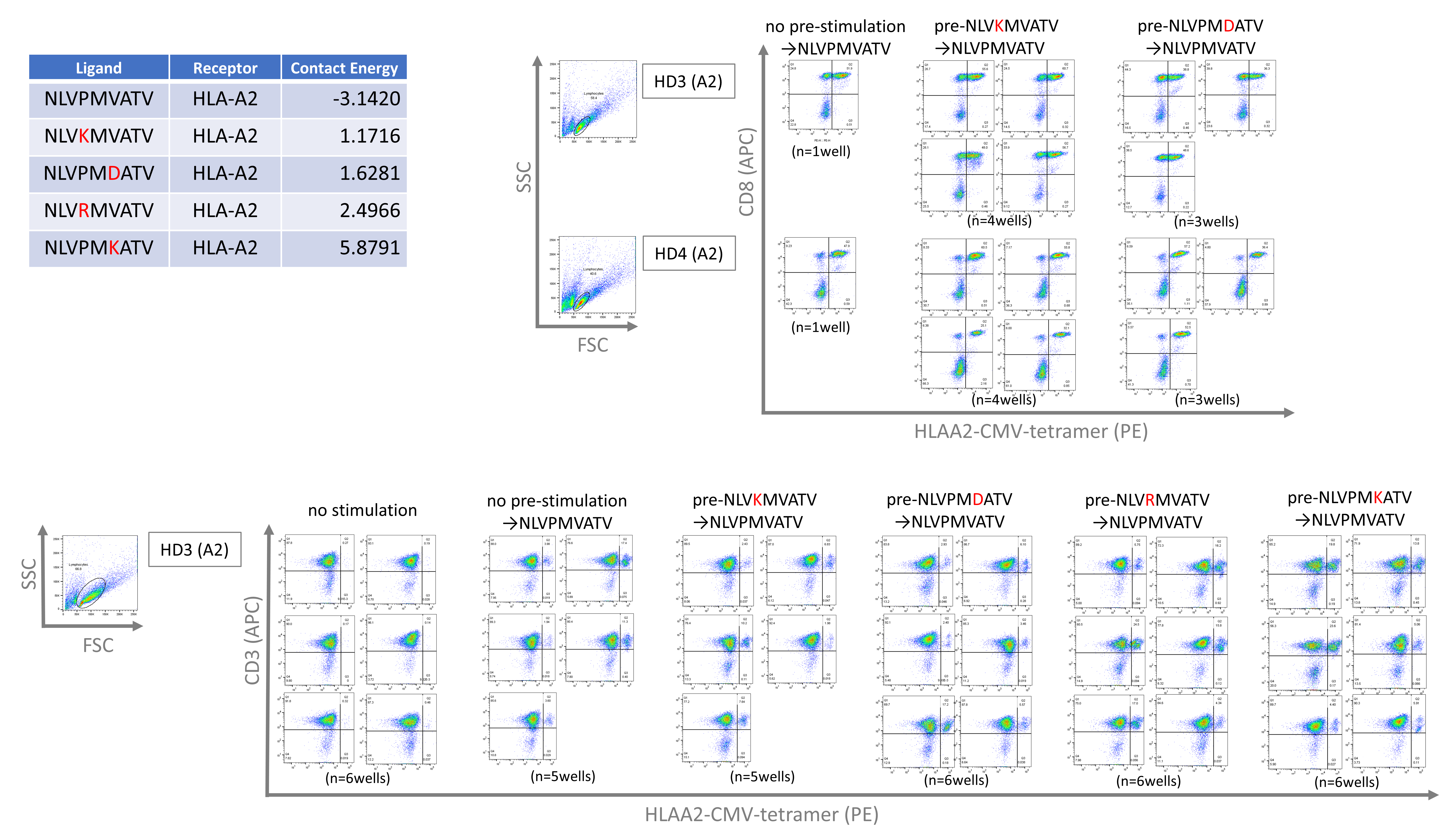

### Fig. S3

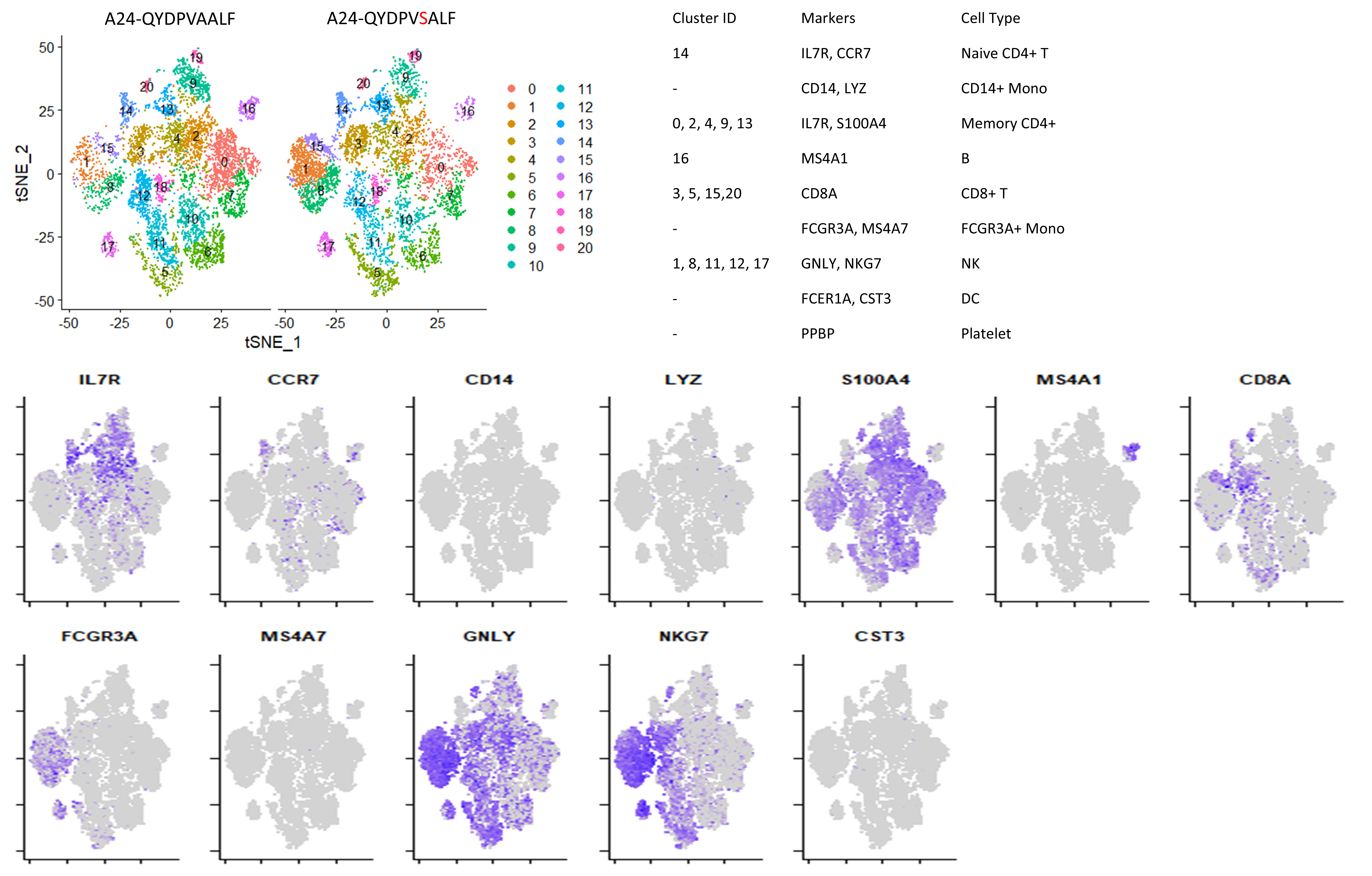

### Fig. S4

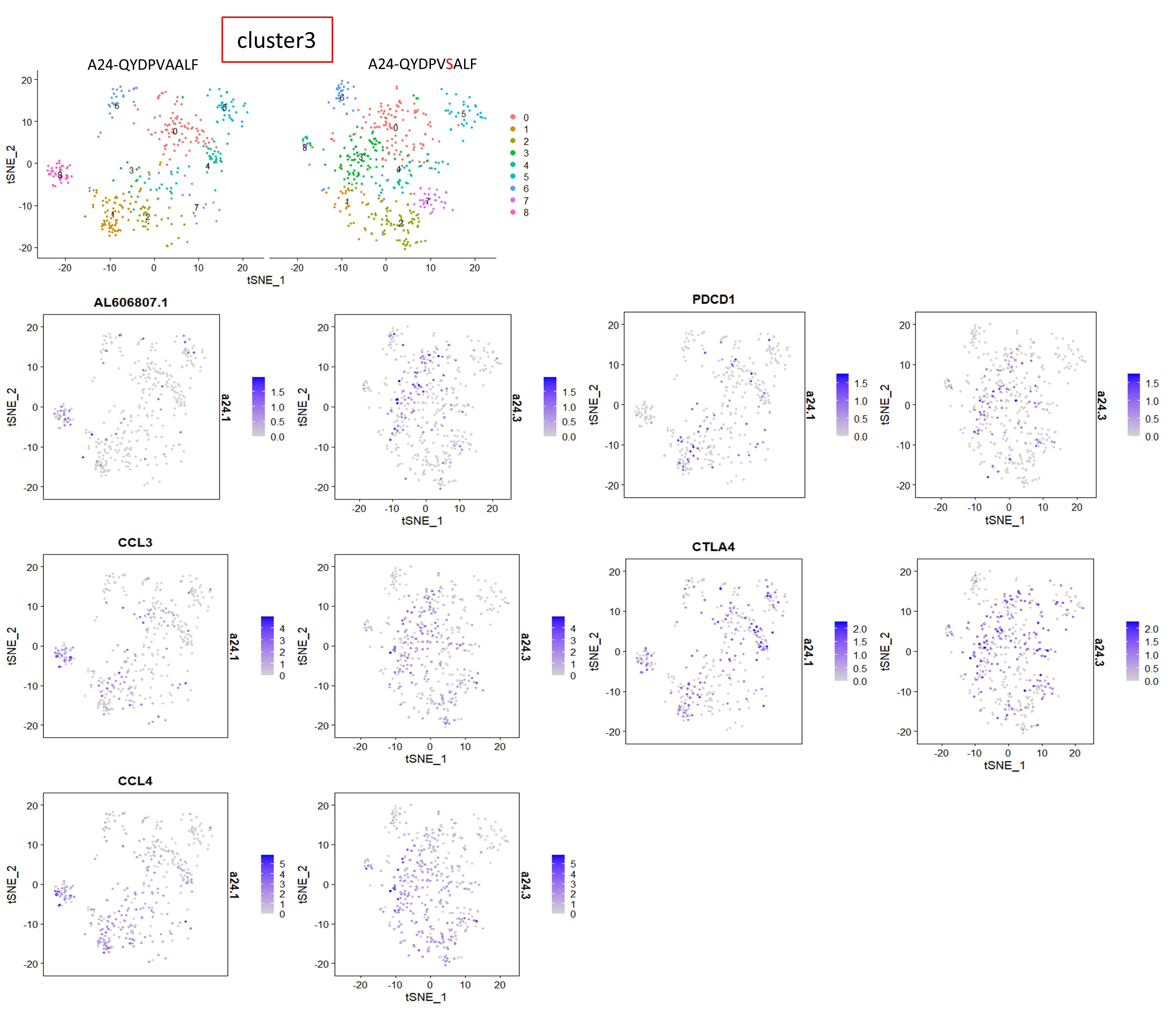

### Fig. S5

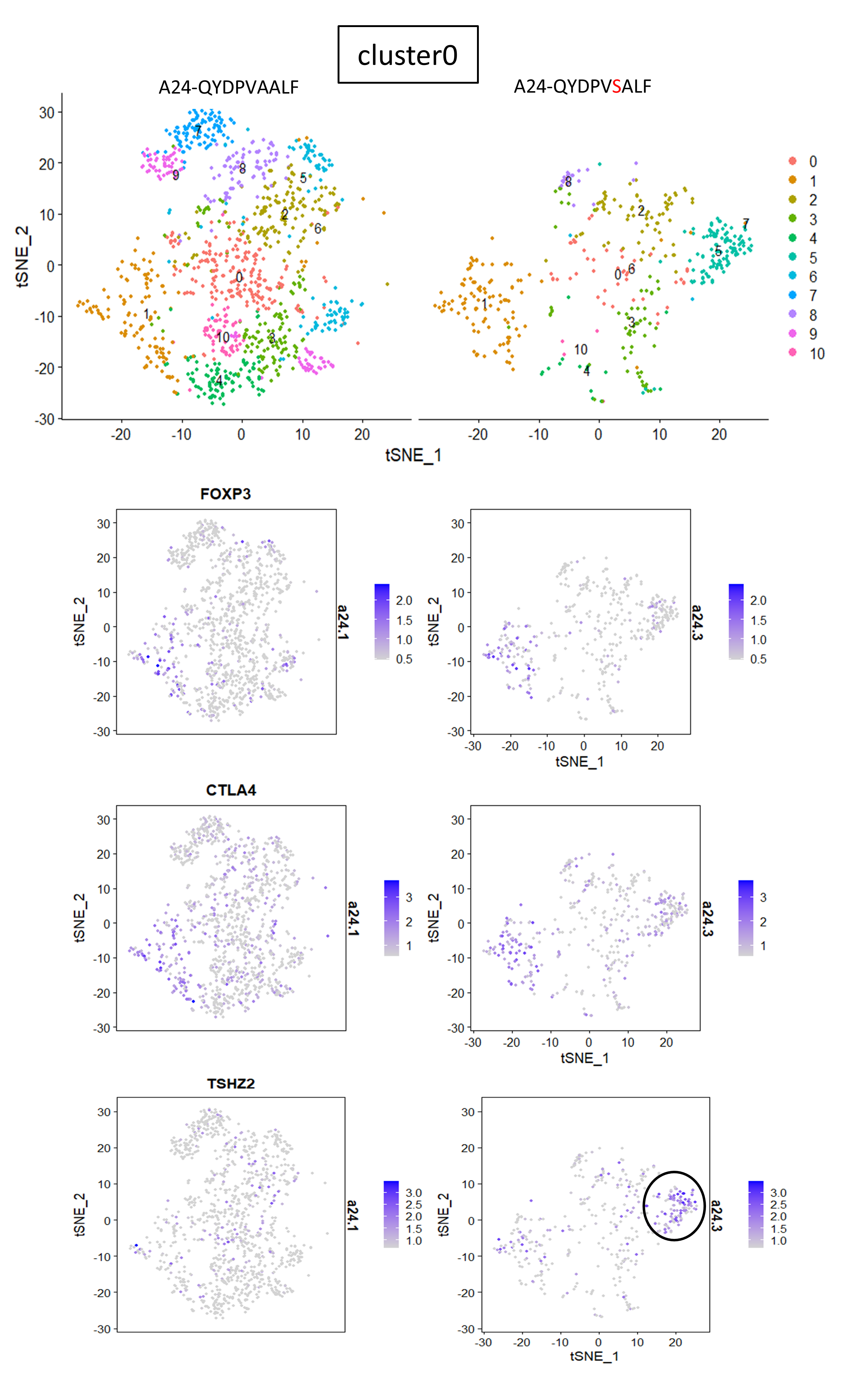
