## Supplementary material for "Induction of anergy in human T lymphocytes by exposure to single amino acid mutated antigens": Table S1

**Table S1** cluster3.3 markers

| | p_val | avg_log2FC |
|---|---|---|
| AL606807.1 | 8.34E-08 | 1.666809 |
| CCL4L2 | 0.004807 | 1.649598 |
| CCL3 | 1.36E-12 | 1.574045 |
| CCL4 | 9.82E-09 | 1.549056 |
| LAYN | 1.60E-07 | 1.442585 |
| PHLDA1 | 2.41E-12 | 1.363277 |
| TNFRSF18 | 4.43E-08 | 1.358842 |
| TBC1D4 | 9.37E-08 | 1.334326 |
| AL138720.1 | 1.28E-11 | 1.247982 |
| AP1S1 | 1.47E-06 | 1.203455 |
| ATP7A | 8.08E-08 | 1.181202 |
| AL136962.1 | 1.01E-05 | 1.180651 |
| TRGV10 | 2.12E-05 | 1.163663 |
| NDFIP2 | 1.84E-12 | 1.159383 |
| PDCD1 | 1.32E-05 | 1.144057 |
