## Supplementary material for "Induction of anergy in human T lymphocytes by exposure to single amino acid mutated antigens": Table S2

**Table S2** cluster 0.5 markers

| | p_val | avg_log2FC |
|---|---|---|
| TSHZ2 | 9.65E-51 | 1.695891 |
| PDE4D | 8.78E-29 | 1.31277 |
| PLPP1 | 9.17E-11 | 1.133 |
| DISC1FP1 | 8.80E-32 | 0.989545 |
| IL4R | 7.68E-08 | 0.921337 |
| TRBV7-9 | 1.99E-113 | 0.894689 |
| SERINC5 | 1.06E-15 | 0.855701 |
| IKZF3 | 3.54E-16 | 0.817792 |
| PTGIS | 2.36E-34 | 0.76632 |
| CD69 | 1.89E-13 | 0.761726 |
| AL109930.1 | 2.38E-33 | 0.739528 |
| PTPRM | 6.82E-30 | 0.733866 |
| LINC01619 | 5.74E-09 | 0.733059 |
| SESN3 | 4.11E-16 | 0.722638 |
| FAM126A | 9.00E-12 | 0.720535 |
